# A meta-analysis of mesophyll conductance to CO_2_ in relation to major abiotic stresses in poplar species

**DOI:** 10.1101/2020.10.19.346270

**Authors:** Raed Elferjani, Lahcen Benomar, Mina Momayyezi, Roberto Tognetti, Ülo Niinemets, Raju Y. Soolanayakanahally, Guillaume Théroux-Rancourt, Tiina Tosens, Mebarek Lamara, Francesco Ripullone, Simon Bilodeau-Gauthier, Mohammed S. Lamhamedi, Carlo Calfapietra

**Author notes:** RE and LB have equally contributed to writing of the manuscript and are joint first-authors. **Highlight** Our meta-analysis on the variation of mesophyll conductance under abiotic stresses shows a noticeable response to light gradient, soil moisture and nitrogen availability and a significant relationship with specific leaf area.

## Abstract

Mesophyll conductance (*g*_m_) determines the diffusion of CO_2_ from the substomatal cavities to the site of carboxylation in the chloroplasts and represents a critical limiting factor to photosynthesis. In this study, we evaluated the average effect sizes of different environmental constraints on *g*_m_ in *Populus* spp., a forest tree model. We collected raw data of 815 *A-C*_i_ response curves from 26 datasets to estimate *g*_m_, using a single curve-fitting method to alleviate method-related bias. We performed a meta-analysis to assess the effects of different abiotic stresses on *g*_m_. We found a significant increase in *g*_m_ from the bottom to the top of the canopy that was concomitant with the increase of maximum rate of carboxylation and light-saturated photosynthetic rate (*A*_max_). *g*_m_ was positively associated with increases in soil moisture and nutrient availability, but insensitive to increasing soil copper concentration, and did not vary with atmospheric CO_2_ concentration. Our results showed that *g*_m_ was strongly related to *A*_max_ and to a lesser extent to stomatal conductance (*g*_s_). Also, a negative linear relation was obtained between *g*_m_ and specific leaf area, which may be used to scale-up *g*_m_ within the canopy.

## Introduction

Carbon assimilation of plants is importantly determined by the diffusion efficiency of CO_2_ from the atmosphere to the site of carboxylation. The rate of CO_2_ diffusion is affected by two main diffusion limitations. The first limitation controls the CO_2_ flux from the atmosphere to the sub-stomatal cavities through the stomata and is characterized by stomatal conductance (*g*_s_). The second limitation determines the diffusion of CO_2_ from the substomatal cavities to the sites of carboxylation in the chloroplasts and is characterized by mesophyll conductance (*g*_m_). *g*_m_ is composed of gaseous and liquid phase resistances (Flexas et al., 2008; Evans et al., 2009; Niinemets et al., 2009). CO_2_ diffusion inside the leaves is complex, facing a series of structural barriers coupled with biochemical regulations. It has been shown that *g*_m_ is typically limited by liquid phase conductance both in species with soft mesophytic leaves as well as in species with tough xerophytic leaves (Tomás et al., 2013; Tosens et al., 2012a,b). The liquid phase is a multi-components pathway that involves the mesophyll cell wall thickness and porosity, the plasmalemma, the chloroplast envelope, the chloroplast thickness and the mesophyll surface area exposed to intercellular air spaces per unit of leaf area (Evans et al., 2009; Tosens et al., 2012b; Tomás et al., 2013). After extensive study during the last two decades, *g*_m_ is now widely accepted as a critical limiting factor to photosynthesis, which has to be considered in characterizing plant carbon gain potentials and responses to future climate change (Evans et al., 2009; Niinemets et al., 2009; Niinemets et al., 2011; Flexas et al., 2016).

Mesophyll conductance has been shown to respond to environmental stress and may govern functional plasticity of photosynthesis and plant fitness under limited resources (Galle et al., 2009; Barbour et al., 2010; Buckley and Warren, 2014; Théroux-Rancourt et al., 2015; Flexas et al., 2016; Shrestha et al., 2018). However, recent findings on the response of *g*_m_ to abiotic stress are conflicting and inconclusive, demonstrating the complex nature of *g*_m_ variation (Flexas et al., 2008; Niinemets et al., 2009; Zhou et al., 2014; Shrestha et al., 2018). This suggests that the environmental and species-specific response (acclimation) of *g*_m_ to growth conditions should be considered to predict plant performance in the field. Among the contrasting environmental responses, growth temperature may (Warren, 2008; Silim et al., 2010) or may not (Dillaway and Kruger, 2010; Benomar et al., 2018) affect *g*_m_. Similarly, the increase in soil nitrogen may (Warren 2004; Shrestha et al., 2018; Xu et al., 2020; Zhu et al., 2020) or may not (Bown et al., 2009) stimulate *g*_m_. The magnitude of decrease in *g*_m_ under water stress and low light differed among studies (Warren et al., 2003; Niinemets et al., 2006; Montpied et al., 2009; Bögelein et al., 2012; Tosens et al., 2012a; Zhou et al., 2014; Peguero-Pina et al., 2015; Théroux Rancourt et al., 2015). These discrepancies among studies result in part from (i) the absolute changes in anatomical, morphological (mesophyll structure) and biochemical (aquaporins and carbonic anhydrase) traits controlling *g*_m_, as well as from changes in the relative contribution of these traits (Marchi et al., 2008; Tomás et al., 2013), and from (ii) the level of coordination between *g*_m_, *g*_s_ and leaf-specific hydraulic conductivity (*K*_L_) (Flexas et al., 2013; Théroux-Rancourt et al., 2014; Xiong et al., 2017). Given the complex interplay between different factors controlling *g*_m_, it is important to examine its acclimation at the genus and species level to gain a general insight into the mechanistic basis of changes in *g*_m_.

Five methods exist to estimate *g*_m_: a) chlorophyll fluorescence coupled to gas exchange (Harley et al., 1992), (b) carbon isotope discrimination coupled to gas exchange (initially developed by Evans et al., 1986), c) oxygen isotope discrimination (Barbour et al., 2016), (d) *A*-*C*_i_ curve fitting (Ethier and Livingston, 2004; Sharkey et al., 2007), and (e) 1D modeling of *g*_m_ from leaf structural characteristics (Evans et al., 2009, Tosens et al., 2012b, Tomas et al., 2013). All of these methods are based on assumptions and each one has its limitations (Flexas et al., 2013; Tosens and Laanisto, 2018). The standard deviation of the estimate of *g*_m_ may vary from 10% to 40%, which may limit our understanding of *g*_m_ acclimation to growth conditions, particularly when the variation between treatments or studies is less than the error of estimates (Sun et al., 2014a).

*Populus* spp., model crops in forestry characterized by high yield potential, have been the subject of numerous studies to understand the physiological response to environmental factors but research is still necessary to make assessment of effects sizes and to draw generalizations (Larocque et al., 2013). A general understanding of the CO_2_ pathway through mesophyll and how it is affected by environmental factors would be beneficial in the efforts to (i) accurately predict canopy photosynthesis under different environmental conditions, particularly under warmer and drier climate, and improve global carbon assimilation models and (ii) develop and effectively select more resilient and productive cultivars for wood, bioenergy and bioremediation. Substantial data of *A-C*_i_ response curves in the literature has been used to estimate photosynthetic parameters, not to estimate *g*_m_, and such compiled dataset would provide a basis to make such assessments on the response of *g*_m_ to the environment.

In this study, we compiled 815 *A*-*C*_i_ response curves from 26 datasets of different poplar species and hybrids (Table 1). Published *A-C*_i_ curve-fitting approaches differ broadly regarding the rectangularity of the hyperbola, segmentations of the model of photosynthesis and determination of the transition value of CO_2_ from carboxylation to electron transport (Harley et al., 1992; Ethier and Livingston, 2004; Manter and Kerrigan 2004; Dubois et al., 2007; Sharkey et al., 2007; Pons et al., 2009; Gu et al., 2010). These approaches led to different fitted values (Miao et al., 2009, Sun et al., 2014a). Although *A-C*_i_ curve fitting is unreliable for species with large *g*_m_, it can provide results similar to those obtained from direct measurements for species with medium to low *g*_m_ (Niinemets et al., 2005, 2006; Qiu et al., 2017; Xu et al., 2020). Using the compiled *A-C*_i_ response curves, we performed curve fitting using a single method (Ethier and Livingston, 2004) to alleviate the fitting method bias and to obtain uniformed estimates of *g*_m_, maximum rate of carboxylation (*V*_cmax_) and rate of electron transport (*J*). We further collected related variables like leaf nitrogen content, stomatal conductance and specific leaf area (SLA) when data were available. Our main goal was to find trends in the response of mesophyll conductance to prevalent abiotic stressors and to examine the relationship between *g*_m_ and other leaf physiological and morphological traits. We believe that a meta-analytical approach to analyze the accumulated data on the diffusion of CO_2_ through the mesophyll diffusion pathway in relation to other photosynthesis-related traits provides key insight into physiological and structural controls on mesophyll conductance and into the environmental plasticity of mesophyll conductance. We aim at contributing to the efforts of improving poplar photosynthetic efficiency in poplar breeding programs, and at improving modelling of global carbon assimilation of biomass and bioenergy crops under climate change.

**Table 1.** List of dataset sources used in the meta-analysis

| Author | Journal | *Populus* species or hybrid parents | Number of genotypes | Treatment | Provenance of plant material | Growth Environment | Number of curves |
| --- | --- | --- | --- | --- | --- | --- | --- |
| Attia et al. 2015 | Journal of Experimental Botany | - *P. balsamifera* L.<br>- *P. simonii* Carrière<br>- *P. balsamifera* L. x *P. simonii* Carrière | 3 | Water stress | Canada | Growth chamber | 15 |
| Benomar et al. | Unpublished data | *P. maximowiczii* A.Henry x *P. balsamifera* L. | 2 | Water stress | Canada | Growth chamber | 12 |
| Benomar 2012 | PhD thesis | - *P. maximowiczii* A.Henry x *P. balsamifera* L.<br>- *P. balsamifera* L. x *P. trichocarpa* Torr. & A.Gray | 2 | Spacing & canopy level | Canada | Plantation | 52 |
| Benomar et al. 2019 | Plos one | - *P. maximowiczii* A.Henry x *P. balsamifera* L.<br>- *P. maximowiczii* A.Henry x *P. nigra* L. | 2 | Temperature & nitrogen | Canada | Growth chamber | 23 |
| Borghi et al. 2007 | Environmental and Experimental Botany | *P. x euramericana* (*P. deltoides* W.Bartram x *P. nigra* L.) (clone Adda) | 1 | Copper | Italy | Growth chamber | 21 |
| Borghi et al. 2008 | Environmental and Experimental Botany | - *P. alba* L.<br>- *P. x Canadensis* (*P. nigra* L. x *P. deltoides* W.Bartram) | 2 | Copper | Italy | Growth chamber | 18 |
| Calfapietra et al. 2005 | Environmental pollution | *P. x euramericana* (*P. deltoides* W.Bartram x *P. nigra* L.) | 1 | Nitrogen & ambient CO<sub>2</sub> & canopy level | Italy | Plantation | 60 |
| Castagna et al. 2015 | Environmental Science and Pollution Research | - *P. x canadensis* (*P. nigra* L. x *P. deltoides* W.Bartram)<br>- *P. deltoides* W.Bartram x *P. maximowiczii* A.Henry | 2 | Ozone & cadmium soil contamination | Italy | Greenhouse | 16 |
| Di Baccio et al. 2009 | Environmental and Experimental Botany | *P. x euramericana* (*P. deltoides* W.Bartram x *P. nigra* L.) (clone i-214) | 1 | Zinc soil contamination | Italy | Growth chamber | 12 |
| Elferjani et al. 2016 | Environmental and Experimental Botany | - *P. trichocarpa* Torr. & A.Gray x *P. balsamifera* L. (clone 747215)<br>- *P. balsamifera* L. x *P. maximowiczii* A.Henry (clones 915004 and 915005)<br>- *P. maximowiczii* A.Henry x *P. balsamifera* L. (clone 915319) | 4 | Latitudinal gradient | Canada | Plantation | 24 |
| Li et al. 2013 | Tree Physiology | *P. euphratica* Oliv. | 1 | Ground water availability | China | In field under shelter (lysimeter) | 9 |
| Merilo et al. 2010 | Baltic Forestry | - *P. nigra* L.<br>- *P. alba* L. | 2 | Ambient CO<sub>2</sub> (FACE) & nitrogen & canopy level | Italy | Plantation | 104 |
| Niinemets et al. 1998 | Tree Physiology | *P. tremula* L. | 1 | Canopy level | Estonia | Natural forest stands | 14 |
| Ripullone et al. 2003 | Tree Physiology | *P. x euramericana* (*P. deltoides* W.Bartram x *P. nigra* L.) (clone i-214) | 1 | Nitrogen | Italy | Greenhouse | 14 |
| Ryan et al. 2009 | Plant, Cell and Environment | *P. deltoides* W.Bartram x *P. trichocarpa* Torr. & A.Gray | 2 | Ozone | United Kingdom | Greenhouse | 118 |
| Silim et al. 2010 | Photosynthetic Research | *P. balsamifera* L. | 1 | Habitat & growth temperature | Canada | Greenhouse | 30 |
| Soolanayakanahally et al. 2009 | Plant, Cell and Environment | *P. balsamifera* L. | 1 | Latitudinal gradient | Canada | Greenhouse | 72 |
| Théroux-Rancourt et al. | Unpublished data | - *P. deltoides* W.Bartram × *P. nigra* L. (clone 3570)<br>- *P. maximowiczii* A.Henry × (*P. deltoides* W.Bartram × *P. trichocarpa* Torr. & A.Gray) (clones 505372 and 505508)<br>- *P. maximowiczii* A.Henry × *P. trichocarpa* Torr. & A.Gray (clone 750361)<br>- *P. maximowiczii* A.Henry × *P. balsamifera* L. (clones 915302, 915313, 915318)<br>- (*P. deltoides* W.Bartram × *P. nigra* L.) × *P. trichocarpa* Torr. & A.Gray (clone 915508) | 8 | N/A | Canada | Greenhouse | 38 |
| Théroux-Rancourt et al. 2014 | Journal of Experimental Botany | - Assiniboine: [(*P.* × '*Walker*': *P. deltoides* W.Bartram × *P. x petrowskiana* R.I. Schrod. ex Regel) × male parent unknown]<br>- Okanese [(*P.* × '*Walker*') × *P. x petrowskiana* R.I. Schrod. ex Regel] | 2 | Water stress | Canada | Greenhouse and growth chamber | 3 |
| Théroux-Rancourt et al. 2015 | Tree Physiology | - (*P. maximowiczii* A. Henry) × (*P. deltoides* W. Bartram × *P. trichocarpa* Torr. & A. Gray)<br>- *P. maximowiczii* A.Henry × *P. balsamifera* L.<br>- '*Walker*' [*P. deltoides* W.Bartram × (*P. laurifolia* Ledeb. × *P. nigra* L.)] × *P. deltoides* W.Bartram<br>- '*Walker*' × *P. petrowskyana* Schr.<br>- *P. balsamifera* L. | 5 | Water stress | Canada | Greenhouse and growth chamber | 12 |
| Tissue et al. 2010 | Tree Physiology | *P. deltoides* W.Bartram | 1 | Phosphorous & ambient CO<sub>2</sub> | Australia | Growth chamber | 76 |
| Tognetti et al. | Unpublished data | *P. x euramericana* (*P. nigra* L. × *P. deltoides* W. Bartram) (clone i-214) |  | Zinc soil contamination | Italy | Greenhouse | 24 |
| Tognetti et al. 2004 | Tree Physiology | - *P. deltoides* W.Bartram × *P. maximowiczii* A.Henry<br>- *P. x euramericana* (*P. deltoides* W.Bartram × *P. nigra* L.) (clone i-214) | 2 | Heavy metals | Italy | Greenhouse | 24 |
| Tosens et al. 2012 | Plant, Cell and Environment | *P. tremula* L. | 1 | Light & water stress | Estonia | Growth chamber | 8 |
| Velikova et al. 2011 | Environmental Pollution | *P. nigra* L. | 20 | Nickel soil contamination | Italy | Growth chamber (climate chamber) | 16 |
| Xu et al. 2020 | Tree Physiology | *P. x euramericana* (*P. deltoides* W.Bartram × *P. nigra* L.) (cv. '74/76') | 1 | Nitrogen & ozone | China | Growth chamber | 6 |

## Materials and methods

### Data collection

Data were collected by a web search in Web of Science, Scopus, and Google Scholar using the following key words: (“*Populus*” or “poplar” or “hybrid poplar” or “aspen”) and (“*V*_cmax_” or “maximum rate of electron transport (*J*_max_)” or “mesophyll conductance”). At this step, abstract of every item was checked to confirm the paper is actually about *g*_m_. Then, we looked at the materials and methods section of selected papers where *A*-*C*_i_ response curves of *Populus* spp. were measured.

To get raw data of *A*-*C*_i_ response curves, we contacted the corresponding authors or co-authors of the targeted studies by e-mail and via ResearchGate. We obtained 23 data sets from published studies and three data sets from unpublished studies (Table 1). Collectively, they provided a total of 815 *A*-*C*_i_ response curves.

The total data of 72 genotypes were collected from measurements on plants growing in plantations (5 studies), or under controlled conditions (greenhouse or growth chamber setups; 21 studies) with optimal and stressful conditions (Table 1). After compiling all *A*-*C*_i_ curves, the quality of the data was assessed based on the following criteria: 1) only curves with at least 2 points in the saturation region (*J* region) were retained; 2) only curves with *P*<0.05 for the p*-*value of fitted curves using the method of Ethier and Livingston (2004) were retained; Consequently, 65 curves that did not meet these conditions were removed; and 3) based on the literature, *g*_m_ values in *Populus* spp. using at least two methods simultaneously never exceeded 1 mol m^−2^ s^−1^ (Singsaas et al., 2004; Flexas et al., 2008; Velikova et al., 2011; Tosens et al., 2012a; Théroux-Rancourt et al., 2014; Momayyezi and Guy, 2017; Xu et al., 2020). Then, *g*_m_ > 1 mol m^−2^ s^−1^ were considered as non-available data (94 entries), and *V*_cmax_ and *J* values retained for further analyses. Furthermore, fewer data point and/or large interval in the intracellular concentration of CO_2_ (*C*_i_ μmol mol^−1^) in the RuBisCO limited phase region are the major source of error and probability of failing *g*_m_ (reached the set bound during the grid search) frequently encountered during *A-C*_i_ curve fitting (Warren, 2006; Miao et al., 2009, Sun et al., 2014a; Moualeu-Ngangue et al., 2016).

### Data subsets

To examine the effect of a given abiotic factor on *g*_m_, we estimated that a minimum of three studies is necessary to have reliable conclusions, regardless of the genotype used. Then, we could come up with subsets of data that focused on the same variable and performed analyses on them separately (identified in the column ‘treatment’ in Table 1). Our first goal was to examine the effect of variations in these factors on *g_m_*, light-saturated photosynthetic rate (*A*_max_), *g*_s_, *J*, *V*_cmax_ and in a second step, the relationships between *g*_m_ and other photosynthetic characteristics (*A*_max_, *g*_s_, *J*, *V*_cmax_). The data subsets included the following environmental factors:

– Canopy level: four studies addressed the photosynthetic activity of leaves at the bottom, middle and top of trees (Niinemets et al., 1998; Merilo et al., 2010; Calfapietra et al., 2005; Benomar, 2012).
– Ambient CO_2_: we examined the response of trees to elevated ambient CO_2_ from Calfapietra et al. (2005), Merilo et al. (2010) and Tissue et al. (2010). We considered 370 ppm as the control treatment in the three studies, while the elevated CO_2_ was 550 ppm of CO_2_ for the studies of Calfapietra et al. (2005) and Merilo et al. (2010), and 700 ppm for the study of Tissue et al. (2010).
– Copper stress: data sets from studies of Borghi et al. (2007) and Borghi et al. (2008) were used to examine the response of poplar trees to contamination of the substrate with copper (Cu). Treatments were assigned to three levels of Cu: 0 (0 to 0.4 μM), 20 (20 to 25 μM) and 75 (75 to 100 μM).
– Soil nitrogen (N) content: High *vs*. low soil N content treatments were reported in four studies: Benomar et al. (2018), Ripullone et al. (2003), Calfapietra et al. (2005) and Xu et al. (2020). In Merilo et al. (2010), no effect of nitrogen fertilization was observed by authors due to high background nutrient availability in the plantation site.
– Soil moisture: water status of trees was assessed and data from four studies was classified into two treatments: control (optimal watering) *vs*. water deficit (Tosens et al., 2012a; Li et al., 2012; Théroux-Rancourt, unpublished; Benomar et al., unpublished).

For Xu et al. (2020), we extracted data from the article (means and standard errors) and generated three replicates assuming a normal distribution using SURVEYSELECT procedure of SAS (SAS Institute, software version 9.4, Cary, NC, USA). The reason is that the authors used the same curve fitting approach (Ethier and Livingston 2004) we used in the meta-analysis (Table 1).

For studies with two or more investigated factors, we considered the different levels of the factor of interest and the control level of the rest of factors to avoid between-factors interaction effects on results. For example, in Calfapietra et al. (2005), trees were subject to different levels of N and CO_2_. When we focused on the effect of N, we selected trees exposed to ambient CO_2_ only (control).

### Curve analysis

Mesophyll conductance and photosynthetic capacity variables, *V*_cmax_ and *J*, were estimated by fitting *A*-*C*_i_ curve with the non-rectangular hyperbola version of the biochemical model of C_3_ plants (Farquhar et al., 1980). The model was fitted using non-linear regression techniques (Proc NLIN, SAS) following Dubois et al. (2007) and Sun et al. (2014a).

Briefly, the net assimilation rate (*A*_n_) is given as:

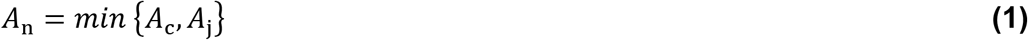

with

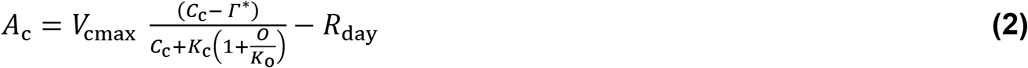

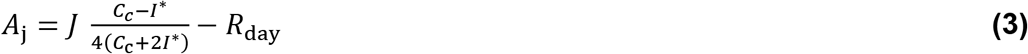

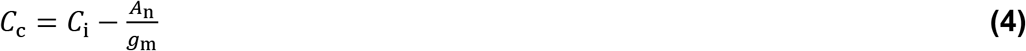

where *V*_cmax_ is the maximum rate of carboxylation (*μ*mol m^−2^ s^−1^), *O* is the partial atmospheric pressure of O_2_ (mmol mol^−1^), *Γ^*^* is the CO_2_ compensation point in the absence of mitochondrial respiration, *R*_day_ is mitochondrial respiration in the light (*μ*mol CO_2_ m^−2^ s^−1^), *C*_c_ is the chloroplast CO_2_ (μmol mol^−1^), *C*_i_ is the intercellular air space concentration of CO_2_ (μmol mol^−1^), *K*_c_ (μmol mol^−1^) and *K*_o_ (mmol mol^−1^) are the Michaelis–Menten constants of RuBisCO for CO_2_ and O_2,_ respectively, *J* is the rate of electron transport (μmol m^−2^ s^−1^). The values at 25 °C used for *K*_c_, *K*_o_ and *Γ^*^* were 272 μmol mol^−1^, 166 mmol mol^−1^ and 37.4 μmol mol^−1^, respectively (Sharkey et al., 2007) and their temperature dependencies were as in Sharkey et al. (2007).

In four data sets, measurements were carried out under a temperature that was different from the reference (25 °C). In this case, *V*_cmax_ and J were normalized to 25 °C using the model of Kattge and Knorr (2007), which integrates the acclimation to growth temperature. However, the actual values of *V*_cmax_ and *J* were more often significant compared to normalized values, and this was true using both ANOVA and regression.

### Statistical analyses

Data analysis assessing the effect of the environmental factors on *g*_m_ and the relationship between *g*_m_ and the other traits were carried out using SAS software (SAS Institute, software version 9.4, Cary, NC, USA).

When at least three studies focused on one factor (nitrogen, CO_2_, canopy level, copper), the effect of treatments on light-saturated photosynthetic rate (*A*_max_), *g*_m_, and *g*_s_ was assessed, separately for each response variable, through mixed model analyses of variance using the primary data (Riley et al., 2010; Mengersen et al., 2013). “Treatment” was the fixed effect while “study” and “genotype” nested within study were the random effects. The number of replicates was not necessarily balanced across treatments. The assumptions of normality of the residuals and homogeneity of variance were verified, and a log-transformation was made when necessary.

## Results

The number of studies on mesophyll conductance has rapidly increased since 2000, and more remarkably since 2013 (Fig. 1), suggesting a growing interest among plant ecophysiologists to understand the role of *g*_m_ in photosynthesis. This pattern was very similar to the increase of publication number on mesophyll conductance in *Populus* spp. (Fig. 2).

**Figure 1.**
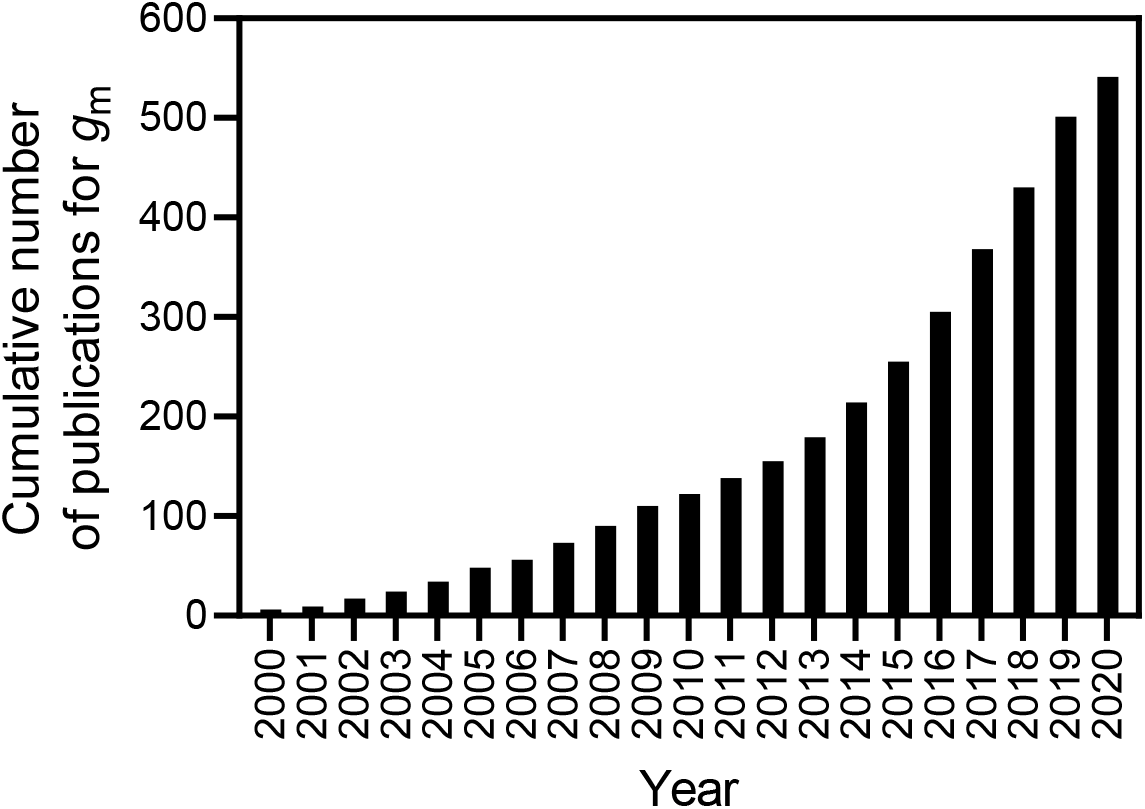
Cumulative number of published studies for mesophyll conductance (*g*_m_) between the years 2000 to 2020. Number of publications were determined using keywords (e.g. *g*_m_) through database search available at the Web of Science Core Collection. (https://clarivate.com/webofsciencegroup/solutions/web-of-science-core-collection/).

**Figure 2.**
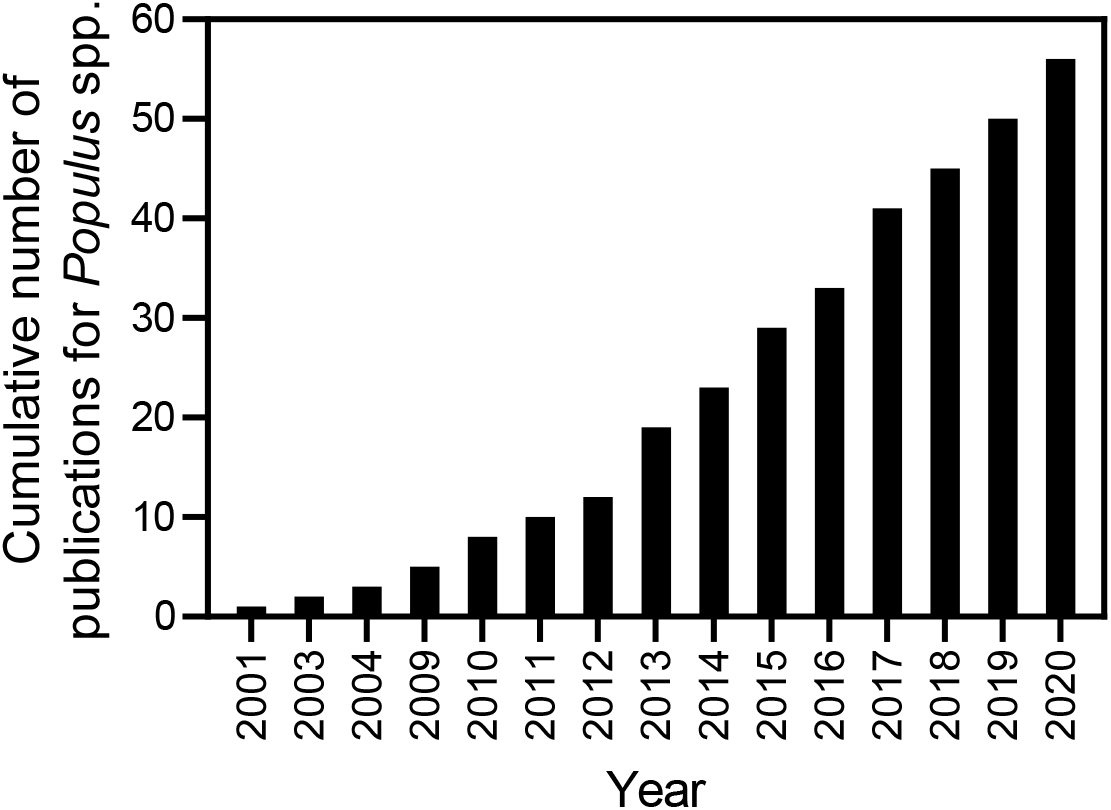
Cumulative number of published studies for mesophyll conductance (*g*_m_) in *Populus* spp. between the years 2001 to 2020. Number of publications were determined using keywords (e.g. *Populus*) through database search available at the Web of Science Core Collection. (https://clarivate.com/webofsciencegroup/solutions/web-of-science-core-collection/).

### Canopy level

Light-saturated photosynthetic rate at an ambient CO_2_ concentration (380-400 μmol mol^−1^), *A*_max_, significantly increased from 7.1±0.44 μmol m^−2^ s^−1^ on average at the bottom leaves to 13.0±0.45 μmol m^−2^ s^−1^ at the mid-canopy, to 16.2±0.53 μmol m^−2^ s^−1^ at the upper canopy (Fig. 3a). Similar to *A*_max_, *g*_m_ had an ascending pattern, from the bottom (0.12±0.01 mol CO_2_ m^−2^ s^−1^) to the top of the canopy (0.24±0.02 mol m^−2^ s^−1^) (Fig. 3c). Stomatal conductance (*g*_s_) was the lowest at the bottom canopy (0.17±0.01 mol H_2_O m^−2^ s^−1^) and then increased to 0.36±0.02 mol H_2_O m^−2^ s^−1^ at the mid and the upper canopy (Fig. 3b). The *g*_m_/*g*_s_ ratio was significantly greater at the upper canopy (1.17±0.1), compared to the mid-canopy leaves (0.88±0.1) and was not different everywhere else (Fig. 3d). While, *V*_cmax_ increased similarly to *A*_max_ and *g*_m_ from the bottom to the top of the canopy (Fig. 3e); however, SLA had an opposite trend (Fig. 3f).

**Figure 3.**
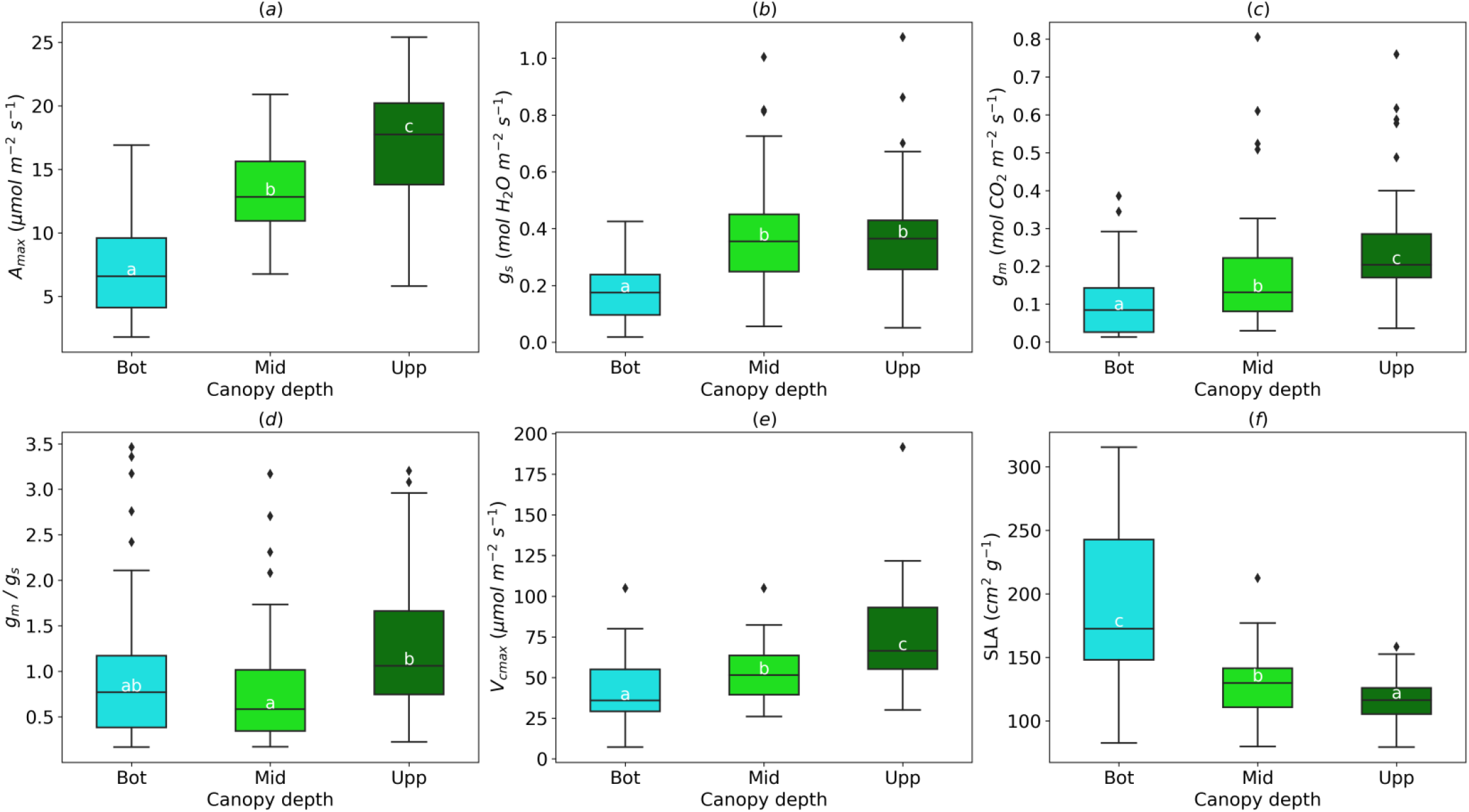
Effect of the leaf position in the canopy (Bottom: Bot, Middle: Mid, Upper: Upp) on light-saturated photosynthetic rate (*A*_max_, a); stomatal conductance (*g*_s_, b); mesophyll conductance (*g*_m_, c); *g*_m_/*g*_s_ ratio (d); maximum rate of carboxylation (*V*_cmax_, e) and specific leaf area (*SLA*, f). In *g*_m_/*g*_s_ ratio, *g*_s_ for water (mol H_2_O m^−2^ s^−1^) was divided by 1.6 to obtain *g*_s_ in mol CO_2_ m^−2^ s^−1^. Means having the same letters are not significantly different at α = 0.05.

### Ambient CO_2_

Increased air CO_2_ had no effect on average *A*_max_ (14.43 ±0.60 μmol m^−2^ s^−1^), *g*_m_ (0.21±0.02 mol m^−2^ s^−1^) and *g*_m_/*g*_s_ (1.09 ±0.1) (Fig. 4a, 4b and 4c). However, average *g*_s_ was higher (0.40 ±0.03 mol H_2_O m^−2^ s^−1^) under ‘‘Ambient’’, compared to ‘‘Elevated’’ CO_**2**_(0.32±0.02 mol H_2_O m^−2^ s^−1^) (Fig. 4b).

**Figure 4.**
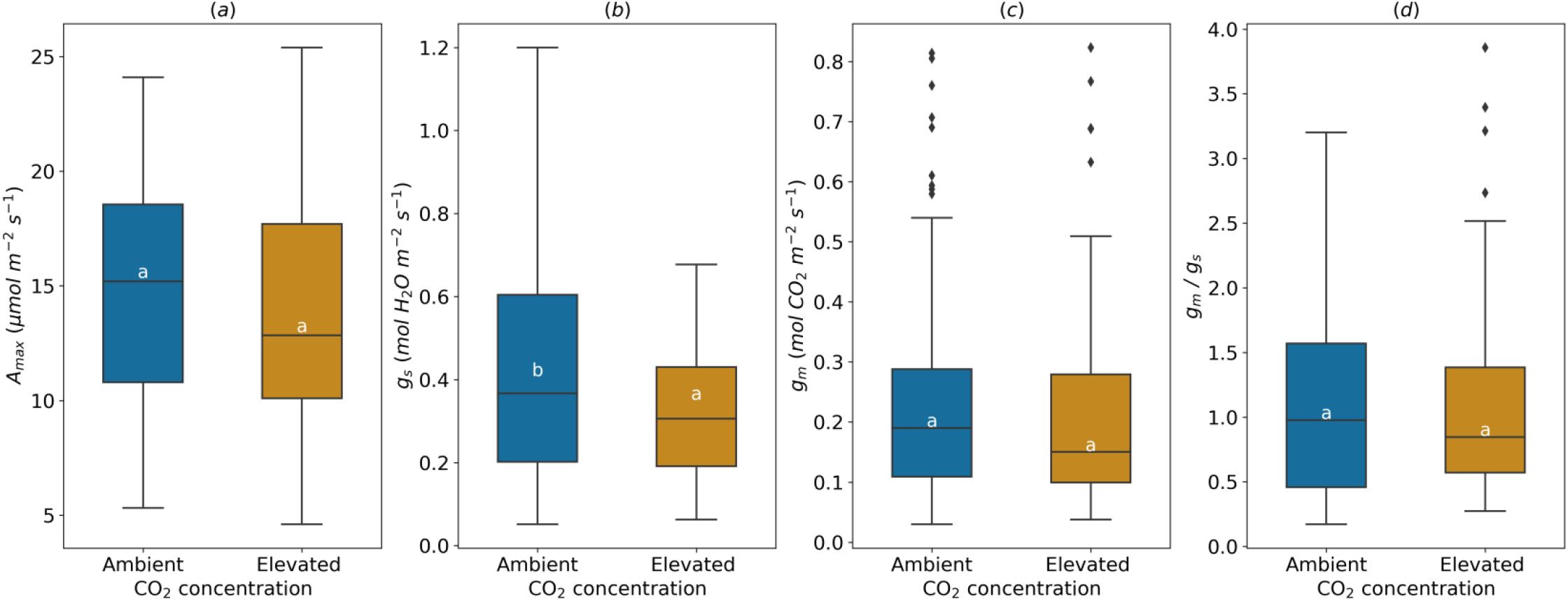
Effect of the ambient air CO_2_ concentration on light-saturated photosynthetic rate (*A*_max_, a); stomatal conductance (*g*_s_, b); mesophyll conductance (*g*_m_, c) and *g*_m_/*g*_s_ ratio (d). In *g*_m_/*g*_s_ ratio, *g*_s_ for water (mol H_2_O m^−2^ s^−1^) was divided by 1.6 to obtain *g*_s_ in mol CO_2_ m^−2^ s^−1^. Means having the same letters are not significantly different at α = 0.05.

### Copper stress

*A*_max_ was not affected when Cu soil concentration increased from 0 to 20 or 75 μM (9.67±0.95 μmol m^−2^ s^−1^) (Fig. 5a). It should be noted that at the highest Cu level (75 μM), *A*_max_ ranged from 4 to 15 μmol m^−2^ s^−1^. Average *g*_s_ significantly decreased under medium (20 μM, 0.17±0.02 mol H_2_O m^−2^ s^−1^), and high Cu treatment (75 μM, 0.18±0.03 mol m^−2^ s^−1^), compared to control treatment (Fig. 5b). Increasing Cu concentration in soil did not affect *g*_m_ (Fig. 5c). The *g*_m_/*g*_s_ ratio was greater under 20 and 75 μM Cu, compared to control (Fig. 5d).

**Figure 5.**
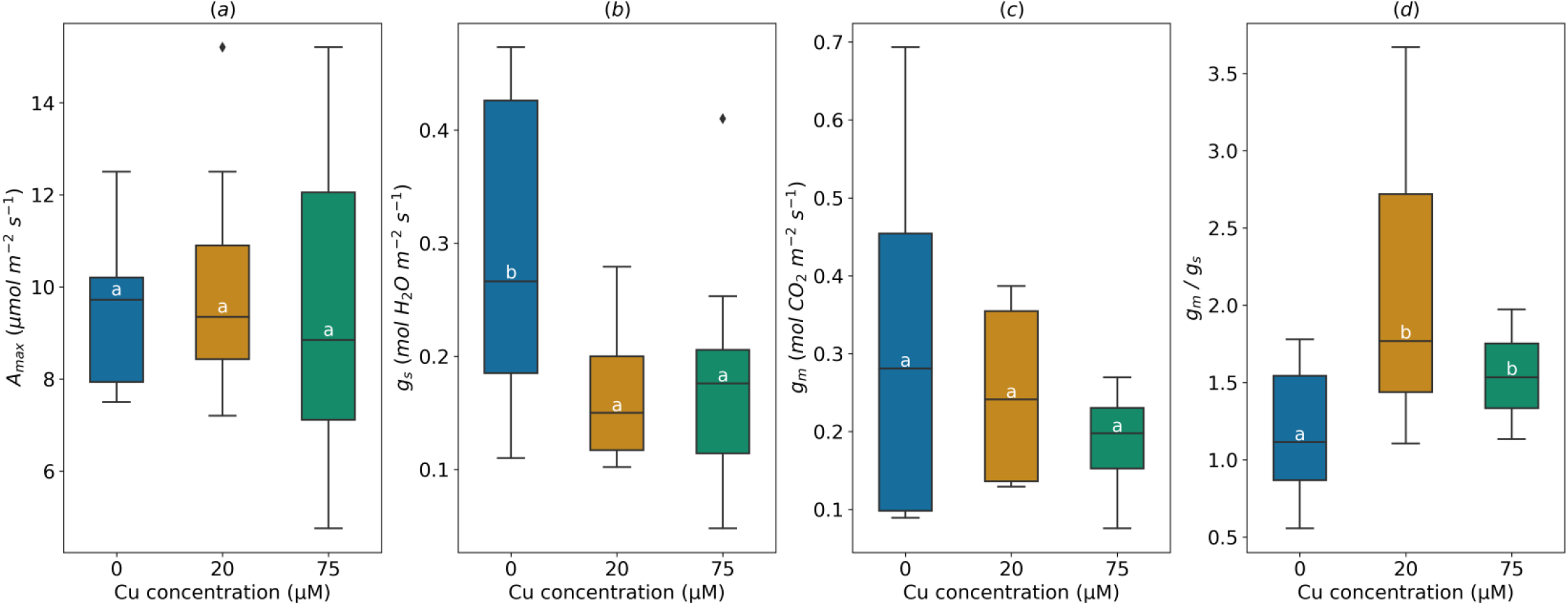
Effect of the soil Copper (Cu) concentration on light-saturated photosynthetic rate (*A*_max_, a); stomatal conductance (*g*_s_, b); mesophyll conductance (*g*_m_, c) and *g*_m_/*g*_s_ ratio (d). In *g*_m_/*g*_s_ ratio, *g*_s_ for water (mol H_2_O m^−2^ s^−1^) was divided by 1.6 to obtain *g*_s_ in mol CO_2_ m^−2^ s^−1^. Means having the same letters are not significantly different at α = 0.05.

### Soil nitrogen

*A*_max_ was significantly greater (16.1 ±0.61 μmol m^−2^ s^−1^) under the high soil nitrogen (HN, 250 kg N ha^−1^ y^−1^ in field study or 20 mM for pot study), compared to the low nitrogen treatment (LN, 12.9±0.65 μmol m^−2^ s^−1^) (Fig. 6a). The high supply of nitrogen increased *g*_s_ (from 0.29±0.03 at LN to 0.36±0.03 mol m^−2^ s^−1^ at HN) and *g*_m_ (from 0.19±0.02 to 0.23±0.02 to mol m^−2^ s^−1^), but had no effect on *g*_m_/*g*_s_ ratio (1.38±0.16 on average) (Fig. 6b, 6c and 6d).

**Figure 6.**
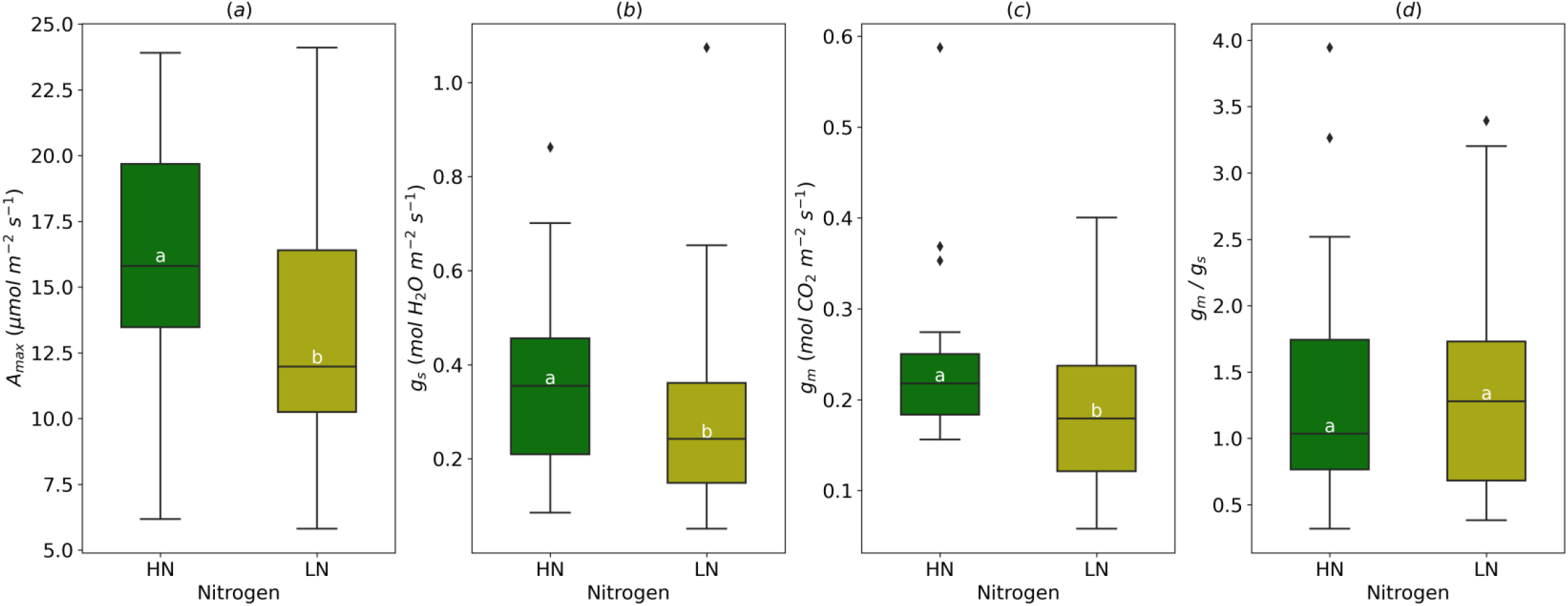
Effect of the soil nitrogen content (high nitogen: HN, low nitrogen: LN) on light-saturated photosynthetic rate (*A*_max_, a); stomatal conductance (*g*_s_, b); mesophyll conductance (*g*_m_, c) and *g*_m_/*g*_s_ ratio (d). In *g*_m_/*g*_s_ ratio, *g*_s_ for water (mol H_2_O m^−2^ s^−1^) was divided by 1.6 to obtain *g*_s_ in mol CO_2_ m^−2^ s^−1^ Means having the same letters are not significantly different at α = 0.05.

### Soil moisture

Average *A*_max_ decreased by drought (range of leaf predawn water potential under water deficit, Ψ_leaf_ = −0.7 to −0.8, soil water content = 10%), dropping from 17.0±0.7 μmol m^−2^ s^−^1 to 14.8± 0.8 μmol m^−2^ s^−1^ on average with minimum value (3.8 μmol m^−2^ s^−1^) much lower than in watered trees (8.9 μmol m^−2^ s^−1^) (Fig. 7a). As expected, soil moisture deficit remarkably altered *g*_s_, decreasing its average value from 0.33±0.02 mol m^−2^ s^−1^ in control trees to 0.20±0.03 mol m^−2^ s^−1^ under drought conditions (Fig. 7b). Drought had the same effect on *g*_m_, but to a lesser extent than *g*_s_. *g*_m_ decreased from 0.27±0.02 to 0.19±0.02 mol m^−2^ s^−1^ under soil moisture deficit (Fig. 7c). In addition, *g*_m_/*g*_s_ ratio increased by 37% when plants were subject to a drought (Fig. 7d).

**Figure 7.**
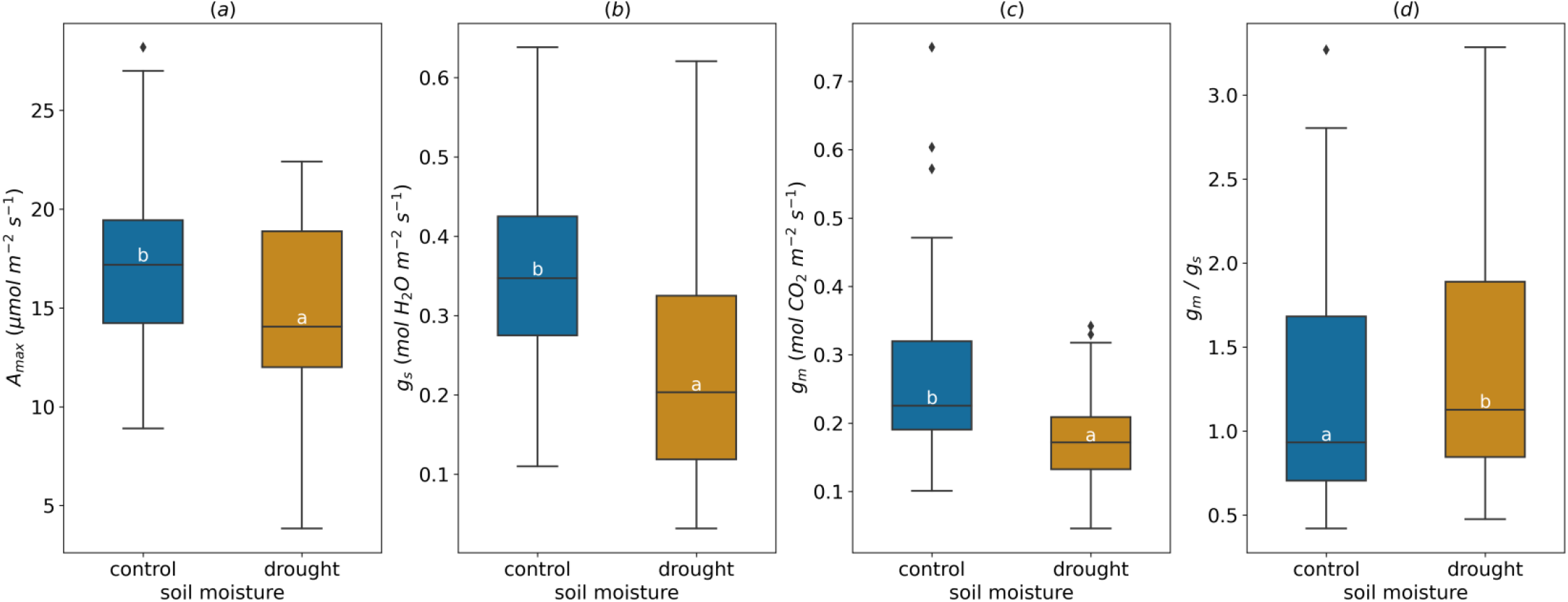
Effect of the soil moisture on light-saturated photosynthetic rate (*A*_max_, a); stomatal conductance (*g*_s_, b); mesophyll conductance (*g*_m_, c) and *g*_m_/*g*_s_ ratio (d). In *g*_m_/*g*_s_ ratio, *g*_s_ for water (mol H_2_O m^−2^ s^−1^) was divided by 1.6 to obtain *g*_s_ in mol CO_2_ m^−2^ s^−1^. Means having the same letters are not significantly different at α = 0.05.

### Relationship between CO_2_ diffusion and photosynthetic activity

*A*_max_ was strongly correlated to both *g*_s_ and *g*_m_ (*P*=0.001) and to *V*_cmax_ (*P*=0.001) over all the studies (Fig. 8a, 8b and 8c). Based on the collected data, *g*_m_ was significantly correlated to *g*_s_ (*P* = 0.04). However, the relationship was not linear. *g*_m_ was the highest (0.4-0.5 mmol m^−2^ s^−1^) when *g*_s_ values were intermediate (0.2-0.4 mol m^−2^ s^−1^), and lowest at high *g*_s_ values (Fig. 8e).

**Figure 8.**
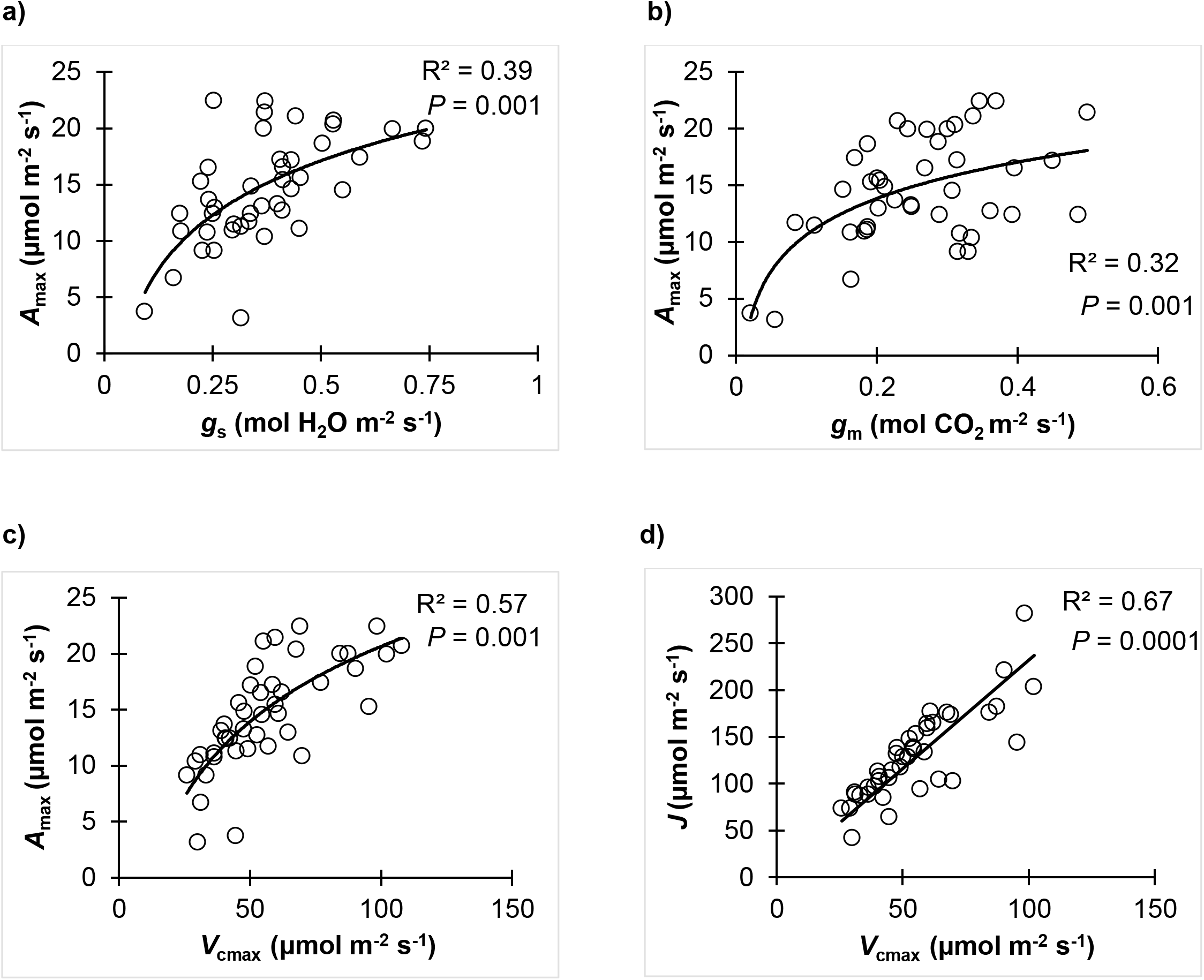
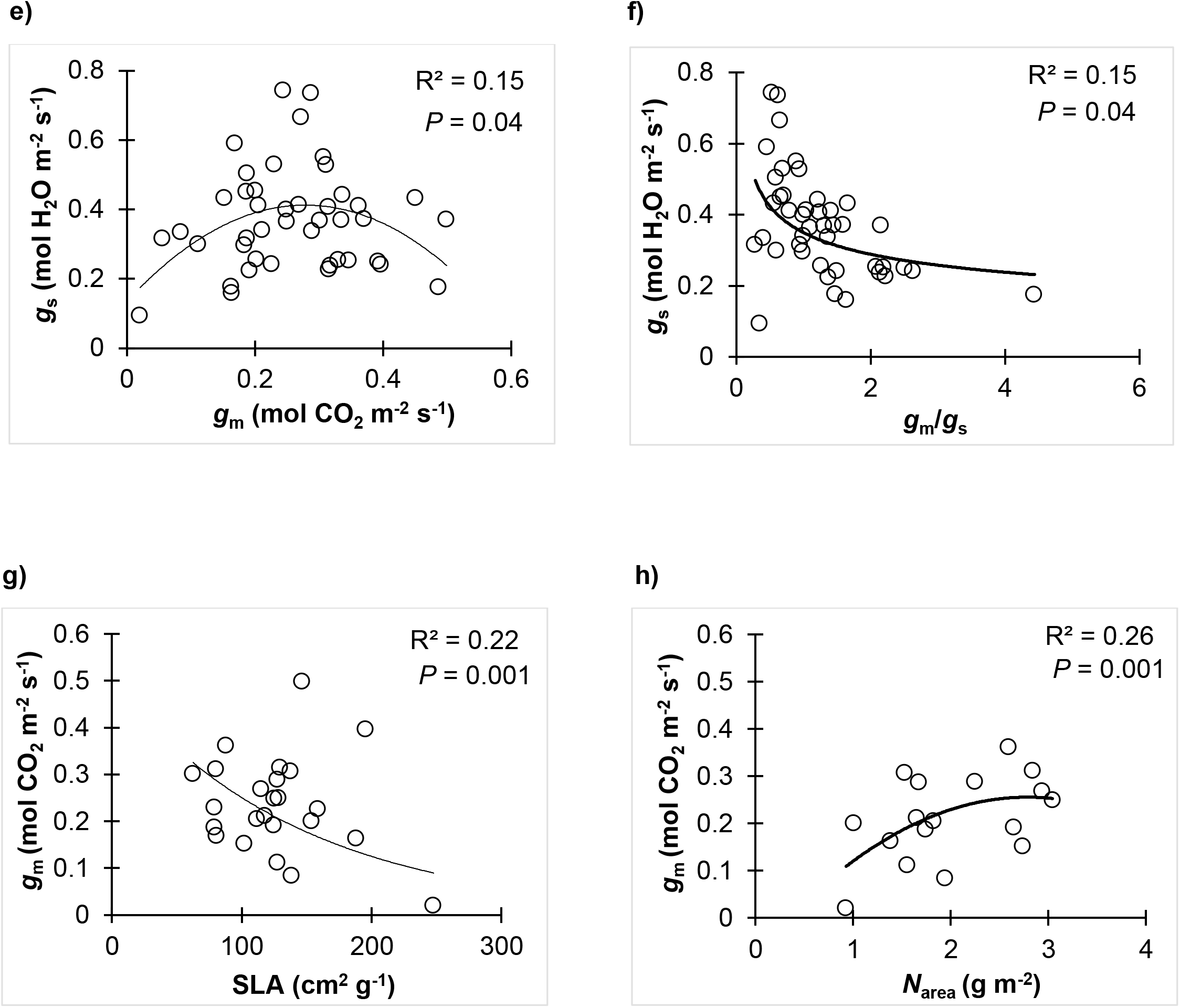
Relationship between light-saturated photosynthetic rate (*A*_max_), stomatal conductance (*g*_s_), mesophyll conductance (*g*_m_), maximum rate of carboxylation (*V*_cmax_), electron transport rate (*J*), *g*_m_/*g*_s_ ratio, specific leaf area (SLA) and per area leaf Nitrogen concentration (*N*_area_). In *g*_m_/*g*_s_ ratio, *g*_s_ for water (mol H_2_O m^−2^ s^−1^) was divided by 1.6 to obtain *g*_s_ in mol CO_2_ m^−2^ s^−1^.

We found a significant negative correlation between SLA and *g*_m_ (*P*=0.001) (Fig. 8g) based on the collected data from studies that measured SLA (n=12). Leaf nitrogen content reported by three studies showed a significant correlation between *g*_m_ and N content per area (*N*_area_) (Fig. 8f). *g*_m_ increased with *N*_area_ until a saturation point (~ 0.25 mol m^−2^ s^−1^).

## Discussion

### Canopy level

The scaling up of photosynthesis from leaves to the canopy and stands (using the model of Farquhar et al., 1980) requires a deep understanding of within-canopy variations in leaf morpho-physiology and the main drivers of foliage acclimation to the dynamic gradient of environmental conditions (light, temperature, vapor pressure deficit (VPD) and soil moisture) (Niinemets et al., 2006; Buckley and Warren, 2014; Niinemets et al., 2015). Unfortunately, pieces of knowledge regarding the variation of *g*_m_ within the canopy and its mechanistic basis are scarce, in particular for *Populus* spp. This situation may explain why most global carbon cycle models remain ‘‘*g*_m_-lacking’’, with possible consequences, such as overestimating the fertilization effect of CO_2_ on global gross primary production and underestimation of water-use efficiency (WUE) and canopy gross photosynthesis under future climate (Sun et al., 2014b; Knauer et al., 2019). The steep and parallel increase of *g*_m_, *A*_max_ and *V*_cmax_ from the bottom to the top of the canopy found here for *Populus* spp. is in agreement with the findings of Niinemets et al. (2006) for *Quercus ilex* L., Montpied et al. (2009) for *Fagus sylvatica* L., and Warren et al. (2003) for *Pseudotsuga menziesii* (Mirbel) Franco. A decrease of *g*_m_ from the bottom to the top of the canopy was also reported (Bögelein et al., 2012; Cano et al., 2013). In absence of water deficit, *g*_m_ limitation is greater under high light (top of the canopy), compared to shade conditions (Niinemets et al., 2009; Cano et al., 2013).

We observed a significant inverse relationship between *g*_m_ and specific leaf area (SLA), comparable to previous studies (Montpied et al., 2009; Niinemets et al., 2006; Tosens et al., 2012b). This suggests that the increase in leaf thickness (lower SLA), e.g. in developing leaves and in leaves grown under higher light, may be associated with increased *g*_m_ (Tosens et al., 2012b). Tosens et al. (2012b) for leaves grown under low and high soil moisture, and Muir et al. (2014) for four species of *Solanum* spp. showed a positive relationship between SLA and *g*_m_. In Tosens et al. (2012b), this relationship reflected increased density of leaves grown under lower water availability. This evidence collectively demonstrates the complex nature of the relationship between SLA and *g*_m_, reflecting the circumstance that SLA is an inverse of the product of leaf thickness and density that can respond differently to environmental drivers (Niinemets, 1999; Poorter et al., 2009). The profile of *g*_m_ within the canopy observed here may be partially attributable to the morphological acclimation of *Populus* spp. foliage to light availability within the canopy. Moreover, this inverse-relationship between SLA and *g*_m_ was used as an empirical model to estimate a maximum attainable *g*_m_ at different canopy layers for C_3_ plant and was implemented in Community Land Model (CLM.4.5) (Sun et al., 2014b; Knauer et al., 2019).

The change in morphological traits and their role in the acclimation of *g*_m_ to a vertical gradient of environmental conditions within the canopy need additional investigations. For instance, shade acclimation of leaf morphology is associated with a lower surface area of chloroplasts exposed to intercellular air spaces (*S*_c_/*S*) and thicker chloroplasts (Hanba et al., 2002; Niinemets et al., 2006; Tosens et al., 2012b; Peguero-Pina et al., 2015). Species-specific leaf development patterns (i.e., evergreen sclerophyllous *vs*. deciduous broadleaves) affect limitations to gas diffusion, thus determining the carbon balance of leaves (Marchi et al., 2007). However, light acclimation may be species-specific and altered by water and soil nitrogen, and leaf ontogeny (Niinemets et al., 2006; Tazoe et al., 2009; Peguero-Pina et al., 2015; Shrestha et al., 2018). It is still unclear whether *g*_m_ profile within the canopy is the result of the change in SLA.

Our results showing higher *g*_s_ and *g*_m_/*g*_s_ at the top of the canopy are in disagreement with the findings of Montpied et al. (2009) and Bögelein et al. (2012), suggesting a species- and environment-specific gradient of *g*_m_/*g*_s_. Temperature and VPD responses of *g*_m_ and *g*_s_ are different (Cano et al., 2013), resulting in different diurnal patterns of *g*_m_ and *g*_s_. Then, the gradient of *g*_m_/*g*_s_ ratio along the canopy may drive the WUE at the canopy level and the midday depression of photosynthetic rate regardless of the level of isohydry of clones (Cano et al., 2013; Buckley and Warren, 2014; Stangl et al., 2019).

### Ambient CO_2_

The response of photosynthetic capacity and diffusion of CO_2_ to free air CO_2_ enrichment (FACE) considerably differed between species and experimental setups. The decrease in *A*_max_ and *g*_s_ in response to elevated CO_2_ showed in our meta-analysis is in agreement with numerous studies on *Populus* spp. and other species (Ainsworth and Rogers, 2007; Medlyn et al., 2013; DaMatta et al., 2016), but is in disagreement with the findings of some other studies, e.g., Sigurdsson et al. (2001) and Uddling et al. (2009). For *g*_m_, the effect of growth CO_2_ changed among studies and some species having an intrinsic low *g*_m_ are more likely to respond to elevated CO_2_ than species with high intrinsic *g*_m_ (Niinemets et al., 2011). However, several studies have reported that *g*_m_ may decrease or be unresponsive to CO_2_ enrichment (Singsaas et al., 2004; Zhu et al., 2012; Kitao et al., 2015; Mizokami et al., 2019). This suggests that the increase of *A*_max_ under elevated CO_2_ cannot be attributed solely to *g*_m_ variation (Singsaas et al., 2004). The absence of *g*_m_ response to elevated CO_2_ complicates the research on mechanisms underlying this variation. Unlike *g*_m_, researchers proposed some hypotheses like least-cost theory, nitrogen limitation and resources investment to explain the decrease of *A*_max_, *V*_cmax_ and *g*_s_ under elevated CO_2_ (Leakey et al., 2009; Smith and Keenan, 2020).

### Copper stress

Similar to our findings, *g*_m_ remained unchanged in the herbaceous plant *Silene paradoxa* L., exposed to high Cu concentration, although *g*_s_ decreased significantly (Bazihizina et al., 2015). In other cases of exposure to other heavy metals, like nickel (Ni), Velikova et al. (2011) reported a significant decrease in chloroplast CO_2_ content and mesophyll conductance in black poplar (*P. nigra* L.) exposed to 200 μM Ni under a hydroponic setup (compared to control = 30 μM Ni). This reduction of *g*_m_ might be attributed to an alteration of leaf structure by toxic effect of high concentrations of heavy metals in mesophyll cells Velikova et al. (2011). Hermle et al. (2007), reported an acceleration of senescence and necrosis of mesophyll cells in *P. tremula* L. leaves exposed to Cu, Zn, Cd and Pb at 640, 3000, 10 and 90 mg per kg of soil, respectively, and a decrease of chloroplast size from the early stages of exposure. The study of Hermle et al. (2007) also reported the thickening of cell walls and change of their chemical composition in damaged mesophyll cells, which might have affected permeability of cell walls and diffusion of CO_2_ through them. Mercury (Hg) (HgCl_2_ form) altered CO_2_ diffusion through aquaporins, a membrane channel of CO_2_ diffusion, in faba bean (*Vicia faba* L.) (Terashima and Ono, 2002) and significantly reduced *g*_m_ in *P. trichocarpa* Torr. & Gray. HgCl_2_ may also decrease *g*_m_ indirectly by disrupting carbonic anhydrase activity, as reported by Momayyezi and Guy (2018), and demonstrated that carbonic anhydrase activity is strongly associated with *g*_m_ variation in *P. trichocarpa* Torr. & Gray (Momayyezi and Guy, 2017).

### Soil Nitrogen

The increase of *A*_max_ by the enhancement of *V*_cmax_ in response to more available soil nitrogen has been established in the literature. However, the possible contribution of *g*_m_ to this augmentation remains unexplored for several species. Our results showed a concomitant increase of *g*_m_ with a higher supply of N. A positive correlation between the level of expression of AQPs genes (PIPs and TIPs) and *g*_m_ has been reported (Hanba et al., 2004; Flexas et al., 2006; Kaldenhoff et al., 2008; Perez-Martin et al. 2014), although it is still unclear whether this is a direct effect or a pleiotropic effect reflecting simultaneous increase in *A*_max_, *g*_m_ and *g*_s_ (Flexas et al., 2012). Recent studies have demonstrated that an increase in *g*_m_ has coincided with an increase in the amount of AQPs after fertilization (Miyazawa et al., 2008b; Zhu et al., 2020).

### Soil moisture

Although many studies showed a decline of *g*_m_ in response to soil water deficit (Flexas et al., 2009; Galle et al., 2009; Tosens et al., 2012a), it remains unclear if this limitation is happening within the mesophyll environment or occurs as a result of a stomatal limitation, which decreases intercellular CO_2_ (*C*_i_). Théroux-Rancourt et al. (2015) showed that, in hybrid poplar, *g*_m_ remained unchanged (~ 0.3 mol m^−2^ s^−1^) following soil drying (Ψ_leaf_ ~ −0.4 to −1.2 MPa), while *g*_s_ decreased, until a threshold of *g*_s_ ~ 0.15 mol m^−2^ s^−1^ from, which *g*_m_ decreased significantly as well. In a trial on *Quercus robur* L. and *Fraxinus angustifolia* Vahl grown in field, Grassi and Magnani (2005) reported a concomitant decrease of both *g*_s_ and *g*_m_ in a dry year (Ψ_soil_ ~ −1.7 MPa), compared with a wetter year (Ψ_soil_ ~ −0.2 MPa). In *P. tremula* L., *g*_m_ significantly declined when Ψ_leaf_ of saplings dropped from −0.3 to −0.7 MPa due to applied osmotic stress (Tosens et al., 2012a). Simultaneously, drought stress induced decrease in SLA accompanied with an increase in the cell wall thickness and a decrease in the chloroplast surface area exposed to intercellular air space per unit leaf area (Tosens et al., 2012a). Other studies have shown that biochemical changes induced by drought stress, like deactivation of aquaporins, could decrease CO_2_ diffusion to carboxylation sites in the chloroplast (Miyazawa et al., 2008a).

Adaptation to the local environment might be a key driver of *g*_m_ variation among taxa, similarly to other morpho-physiological traits. Interspecific and intraspecific differences in *g*_m_ from mesic *versus* xeric environments (*Quercus* spp. and *Eucalyptus* spp.) were reported by Zhou et al. (2014). Their study showed that *g*_m_, as well as *g*_s_, *V*_cmax_ and *J* of species from drier regions were less sensitive to water deficit which maintains transpiration and photosynthesis activity at higher rates under drought, compared to species from the mesic environment. Marchi et al. (2008) observed that structural protection of mesophyll cells had a priority over functional efficiency of photochemical mechanisms in *Olea europaea* L. (evergreen sclerophyllous) but not in *Prunus persica* L. (deciduous broadleaf), depending on age-related variation in mesophyll anatomy.

### Conclusion and Future Directions

The present review shows that *g*_m_ in *Populus* spp. varies predictably along light gradients and that it responds to changes in soil moisture and nutrient availability, but is not affected by metal concentration and increasing atmospheric CO_2_ concentration. Although metabolic processes noticeably influence the response of *g*_m_ to environmental changes, physical constraints through leaf development and ageing need to be considered in scaling photosynthesis from leaf to canopy, and in breeding programs for high WUE. Because fast-growing *Populus* spp. trees are important players in combating climate change, mitigating carbon emissions to some extent, comparisons of genotypes with different adaptations to changing environments and breeding for novel genotype-climate associations are urgently needed. This study shows that the variability of *g*_m_ in different experimental conditions offers a potential indicator for improving *Populus* spp. productivity and resilience. However, more research is yet needed, also combined with anatomical studies, to better understand the sources of variation of CO_2_ diffusion through the mesophyll and their consequences on carbon assimilation and growth.

Moreover, determination of the efficiency and optimal age for early selection of fast-growing poplar clones require an understanding of the genetic control and age-based genetic correlations for traits related to *g*_m_ and growth. For that, a detailed evaluation of the genotypic control of the variances and clonal heritability of *g*_m_ is needed. Finally, the identification of molecular bases of the regulation of *g*_m_ is necessary to further refine a multi-criteria early selection approach of poplar clones dedicated to the future forestry capable of ensuring better productivity and increased resistance to environmental stresses (frost, drought, water logging, heavy metals, heat waves, etc.).

## Acknowledgements

This research was supported by the University of Québec in Abitibi-Témiscamingue (UQAT) as startup funds to ML. The authors acknowledge researchers who kindly provided data used in the meta-analysis.

## Data Availability

The datasets generated for this study are available on request to the corresponding author.

